# Population genomics of the pathogenic yeast *Candida tropicalis* identifies hybrid isolates in environmental samples

**DOI:** 10.1101/2020.11.25.397612

**Authors:** Caoimhe E. O’Brien, João Oliveira-Pacheco, Eoin Ó Cinnéide, Max A. B. Hasse, Chris Todd Hittinger, Thomas R. Rogers, Oscar Zaragoza, Ursula Bond, Geraldine Butler

## Abstract

*Candida tropicalis* is a human pathogen that primarily infects the immunocompromised. Whereas the genome of one isolate, *C. tropicalis* MYA-3404, was originally sequenced in 2009, there have been no large-scale, multi-isolate studies of the genetic and phenotypic diversity of this species. Here, we used whole genome sequencing and phenotyping to characterize 77 isolates *C. tropicalis* isolates from clinical and environmental sources from a variety of locations. We show that most *C. tropicalis* isolates are diploids with approximately 2 - 6 heterozygous variants per kilobase. The genomes are relatively stable, with few aneuploidies. However, we identified one highly homozygous isolate and six isolates of *C. tropicalis* with much higher heterozygosity levels ranging from 36 - 49 heterozygous variants per kilobase. Our analyses show that the heterozygous isolates represent two different hybrid lineages, where the hybrids share one parent (A) with most other *C. tropicalis* isolates, but the second parent (B or C) differs by at least 4% at the genome level. Four of the sequenced isolates descend from an AB hybridization, and two from an AC hybridization. The hybrids are *MTL***a**/*α* heterozygotes. Hybridization, or mating, between different parents is therefore common in the evolutionary history of *C. tropicalis*. The new hybrids were predominantly found in environmental niches, including from soil. Hybridization is therefore unlikely to be associated with virulence. In addition, we used genotype-phenotype correlation and CRISPR-Cas9 editing to identify a genome variant that results in the inability of one isolate to utilize certain branched-chain amino acids as a sole nitrogen source.

**Author summary:** *Candida tropicalis* is an important fungal pathogen, which is particularly common in the Asia-Pacific and Latin America. There is currently very little known about the diversity of genotype and phenotype of *C. tropicalis* isolates. By carrying out a phylogenomic analysis of 77 isolates, we find that *C. tropicalis* genomes range from very homozygous to highly heterozygous. We show that the heterozygous isolates are hybrids, most likely formed by mating between different parents. Unlike other *Candida* species, the hybrids are more common in environmental than in clinical niches, suggesting that for this species, hybridization is not associated with virulence. We also explore the range of phenotypes, and we identify a genomic variant that is required for growth on valine and isoleucine as sole nitrogen sources.

## Introduction

*Candida tropicalis* is an opportunistic pathogenic yeast, and a cause of both superficial and systemic infections in humans. Although *Candida albicans* remains the most common cause of candidiasis, other *Candida* species such as *C. tropicalis* are increasingly isolated as the cause of invasive *Candida* infections [1–3]. *C. tropicalis* is particularly prevalent in Asia-Pacific and Latin America, where it has been identified as the second- or third-most common cause of candidiasis [1–5]. *C. tropicalis* is particularly associated with infection in patients with hematological malignancies [5,6]. Fluconazole and voriconazole resistance occurs more frequently in clinical isolates of *C. tropicalis* than in clinical isolates of *C. albicans* [1,2]; the frequency of resistant isolates, particularly to fluconazole, ranges from 5 - 36% [2,7–10]. Notably, more Asia-Pacific isolates are fluconazole-resistant in comparison to isolates from other locales [1–3]. Bloodstream infections by *C. tropicalis* are associated with high mortality rates, ranging from 41 - 61% [11–13].

*C. tropicalis* is a member of the CUG-Ser1 clade, a group of species in which the CUG codon is translated as serine instead of the standard leucine [14,15]. The genome of *C. tropicalis* was first sequenced in 2009, revealing a diploid genome of approximately 14.5 Mb [16]. Although once thought to be asexual, it is now known that *C. tropicalis* can mate via a parasexual cycle [17,18]. Cells that are homozygous for either the *MTL***a** or *MTLα* mating idiomorph undergo phenotypic switching to the opaque state, and subsequently mate with cells that are homozygous for the opposite mating type [17,19]. The resulting tetraploid heterozygous *MTL***a**/*α* cells undergo concerted chromosome loss to revert to the diploid state [18]. Same-sex mating (i.e. mating between two cells homozygous for the same mating type) has been observed in this species, but only in the presence of the pheromone from the opposite mating type [19]. The majority of *C. tropicalis* isolates (79 - 96%) are heterozygous at the *MTL*, implying that the variation conferred by sexual reproduction is largely beneficial [20,21].

To date, there are no population genomics studies of *C. tropicalis* isolates, although multi-locus sequence typing (MLST) suggests that there is a diverse population structure [22,23]. In contrast, analysis of almost 200 genomes from *C. albicans* isolates identified a clonal population structure with high levels of heterozygosity (e.g. single nucleotide polymorphisms, or SNPs) between the haplotypes of isolates in most lineages [24]. There was also some evidence for gene flow between *C. albicans* lineages [24]. Recent analysis suggests that all isolates of *C. albicans* descended from an ancient hybridization event between related parents, followed by extensive loss of heterozygosity [25].

Some other diploid species from the CUG-Ser1 clade with higher levels of heterozygosity than *C. albicans* also arose from hybridization (or mating) between two related but distinct parents [26–28]. Like *C. albicans*, all currently characterized isolates of *C. metapsilosis* arose from a single hybridization between two unknown parents, followed by rearrangement at the *MTL***a** locus [27]. Similarly, *Millerozyma (Pichia) sorbitophila* is an interspecific hybrid between one parent that is highly similar to *Millerozyma (Pichia) farinosa* and a second unidentified parent which has a high degree of synteny with the first parent, but diverges at the sequence level by about 11% [29]. Hybridization appears to be ongoing in *C. orthopsilosis*, where most isolates descend from one of at least four hybridization events between one known parent with a homozygous genome, and one that differs by about 5% at the genome level [26,28]. In contrast, sequenced isolates of *Candida dubliniensis, Candida parapsilosis* and *C. tropicalis* are not hybrids [25].

Hybridization between two genetically divergent parents is hypothesized to drive adaptation of organisms to new or changing environments. For example, hybridization within the *Saccharomyces* species complex is associated with the development of favorable traits, such as cryotolerance in the lager-brewing yeast *Saccharomyces pastorianus*, a hybrid of *Saccharomyces cerevisiae* and *Saccharomyces eubayanus* [30] or increased thermotolerance and cryotolerance in various hybrids of *S. cerevisiae, S. eubayanus* and *Saccharomyces kudriavzevii* [31]. Other members of the Saccharomycotina are also hybrids, such as the yeast *Zygosaccharomyces rouxii*, used in the production of soy sauce and balsamic vinegar [32]. Some isolates of this species are haploid, while some are highly heterozygous diploids resulting from the hybridization of two parental *Zygosaccharomyces* species [33–35]. The *Cryptococcus neoformans* species complex, which includes several human pathogens, has also been found to include several hybrids, resulting from multiple recent hybridization events between different serotypes [36,37]. Hybridization has been proposed to drive virulence properties, for species within the CUG-Ser1 clade like *C. metapsilosis* [38], and species outside the clade, like *Candida inconspicua* [39].

Here we carried out a population genomic study of 77 *C. tropicalis* isolates, including some from clinical sources and some isolated from the environment. We found that heterozygosity levels range from 2 to 6 variants per kilobase in most isolates. However, one isolate is very homozygous, and six isolates have very heterozygous genomes. The heterozygous isolates appear to be the product of hybridization between one parent that is similar to the *C. tropicalis* reference strain MYA-3404, and other parents that differ from the reference strain by 4 - 4.5%. The hybrid isolates were predominately found in environmental niches, suggesting that hybridization in this species is not associated with virulence. In addition, we characterized the growth phenotypes of the non-hybrid isolates in different environmental conditions, and we associated phenotypic variation with genotypic variation. We found that a deletion of two bases in the gene *BAT22* is associated with the inability of three different *C. tropicalis* strains to use valine and isoleucine as sole nitrogen sources.

## Results

### Population study of *C. tropicalis*

The original reference genome sequence of *C. tropicalis* MYA-3404 was sequenced in 2009, resulting in a genome assembly consisting of 23 supercontigs totaling 14.6 Mb with 6,258 annotated genes [16]. We used Illumina data from resequencing of the reference strain to assemble the 23 supercontigs into 16 scaffolds, called Assembly B (see Materials & Methods). The assembly was subsequently further improved as described by Guin et al [40].

77 unique *C. tropicalis* isolates from different geographical locations were collected and sequenced using Illumina technology. For convenience, we named these strains ct01 to ct78, including only one of two isolates with very similar sequences (Table S1). Most isolates came from clinical sources from the USA, Spain and Ireland. Twelve environmental isolates were included, eleven collected from soil in the USA and Ireland, and one from coconut water in India. The reference strain *C. tropicalis* MYA-3404 (ct11), which was previously sequenced by Sanger sequencing [16], was also resequenced, as were three engineered auxotrophic derivatives in two genetic backgrounds [41,42].

Variants were identified by mapping reads to *C. tropicalis* MYA-3404 Assembly B and calling variants with the Genome Analysis Toolkit (GATK) [43]. Analysis of the distribution of allele frequencies in heterozygous biallelic SNPs showed that the majority of isolates are diploid, i.e. the ratio of reference to non-reference allele frequency is 50:50. However one isolate, *C. tropicalis* ct66 is triploid (peaks of allele frequency at 0.33 and 0.66), and another isolate, *C. tropicalis* ct26, appears to be octaploid (peaks of allele frequency at approximately 0.5, 0.12 and 0.87) (Fig. S1). In addition, we observed single-chromosome aneuploidies in three isolates (Fig. S1). *C. tropicalis* ct06 and *C. tropicalis* ct18 each have three copies of scaffold 8, and *C. tropicalis* ct15 has three copies of scaffold 4 (trisomy). *C. tropicalis* ct15 (CAY3763, derived from *C. tropicalis* AM2005/0093) was used as the background to generate gene deletions [42], a process that has been found to commonly induce aneuploidies in *C. albicans* [44].

Most isolates have approximately 2 - 6 heterozygous variants per kilobase similar to the type strain [16] (Fig. 1A). This is comparable to the level of heterozygosity seen in *C. albicans* (2.5 - 8.6 SNPs per kilobase) [26 [16]]. One isolate (*C. tropicalis* ct20) is extremely homozygous, with 0.84 heterozygous variants per kilobase. This isolate also has a higher proportion of homozygous variants compared to the reference (83% of total variants are homozygous, compared to an average of 41% in other isolates). However, six isolates have exceptionally high levels of heterozygosity (Fig. 1A). These include one clinical isolate from Spain (*C. tropicalis* ct25), and five environmental isolates from soil, one from the USA (*C. tropicalis* ct42) and four from Ireland (*C. tropicalis* ct75, ct76, ct77 and ct78). These isolates have 36 - 49 heterozygous variants per kilobase. Phylogenetic analysis shows that most isolates cluster together (Cluster A in Fig. 1B). However, the six heterozygous isolates are extremely divergent (Cluster B, Fig. 1B). These six isolates separate into two groups, one containing *C. tropicalis* ct25, ct42, ct75 and ct76, and a second containing *C. tropicalis* ct77 and ct78.

**Figure 1.**
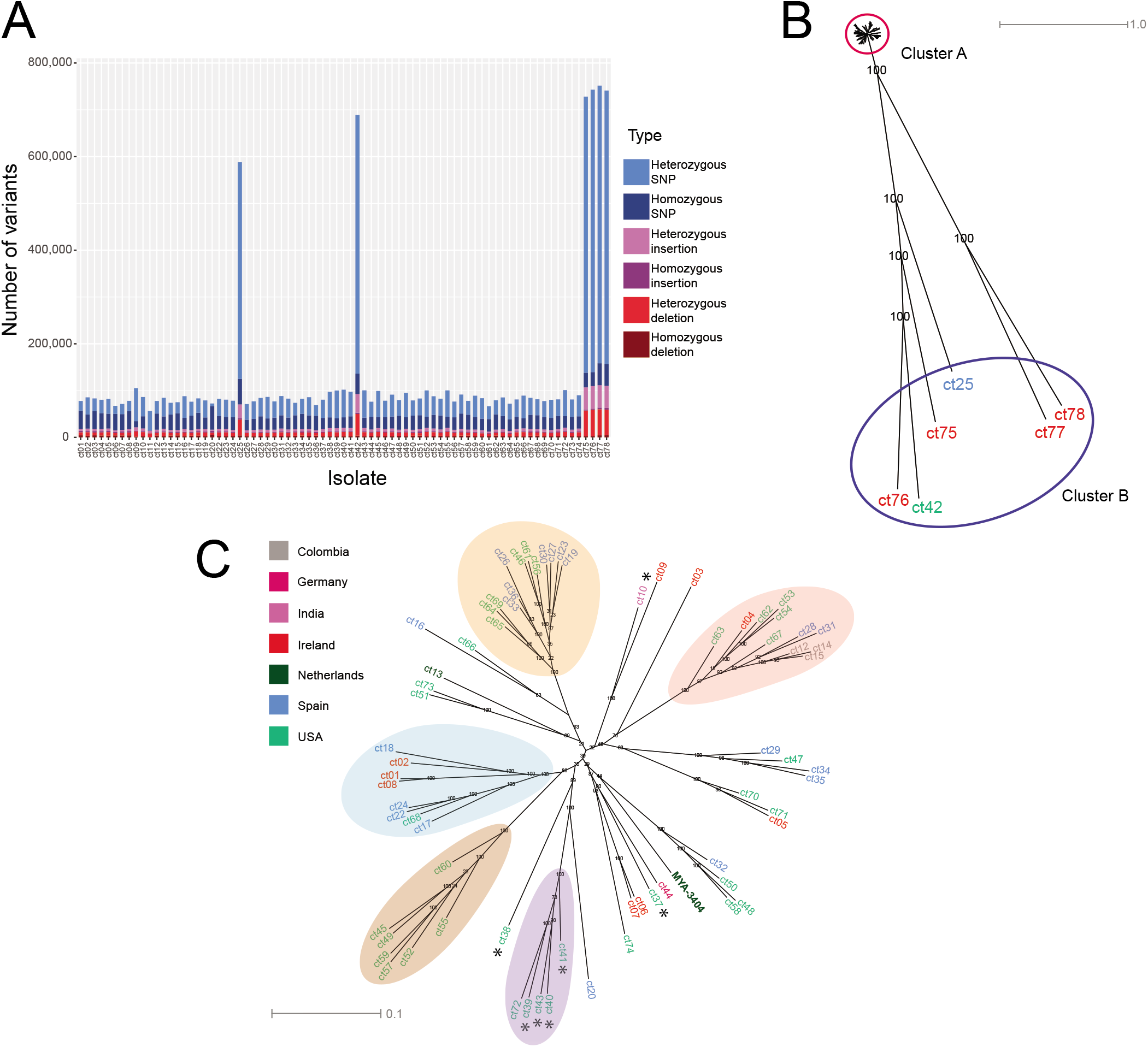
Identification of novel isolates of *C. tropicalis*. **(A) Genome variation among *C. tropicalis* isolates.** Variants were identified using the Genome Analysis Toolkit HaplotypeCaller and filtered based on genotype quality (GQ) scores and read depth (DP). Variants for all 77 isolates are shown according to variant type. Isolates are labelled on the X-axis by strain ID. One isolate (*C. tropicalis* ct20) has mostly homozygous variants, and six isolates have very high levels of heterozygous variants. **(B) Six isolates of *C. tropicalis* are highly divergent.** Variants were called as in (A). For heterozygous SNPs, a single allele was randomly chosen using RRHS [65] and for homozygous SNPs, the alternate allele to the reference was chosen by default. This process was repeated 100 times and 100 SNP trees were drawn with RAxML using the GTRGAMMA model [66]. The best-scoring maximum likelihood tree was chosen as a reference tree and the remaining 99 trees were used as pseudo-bootstrap trees to generate a supertree. Pseudo-bootstrap values are shown as branch labels. The six divergent isolates (Cluster B) are labelled according to their country of origin (see 1C). **(C) SNP phylogeny of isolates from Cluster A indicates that clade structure is not associated with geography**. The phylogeny of cluster A is shown in detail. Pseudo-bootstrap values are shown as branch labels. Isolates are labelled according to their country of origin, and environmental isolates are indicated with an asterisk. The reference strain, *C. tropicalis* MYA-3404, is labelled. The five colored clades are mostly supported by principal component analysis (Fig. S2).

The remaining isolates (Cluster A) are shown in more detail in Fig. 1C. There is evidence of some population structure, with at least five well-supported clades (colored ovals in Fig. 1C, Table S4) and many lineages outside these clades. However, there is little obvious correlation between phylogeny and geography. Two clades contain only isolates from the USA, but this likely reflects the overrepresentation of isolates from the USA in our collection. In addition, although some of the environmental isolates cluster together, others are closely related to clinical isolates (Fig. 1C). There is therefore no clear distinction between clinical and environmental isolates.

### Origins of the heterozygous *C. tropicalis* isolates

The levels of heterozygosity in the six divergent *C. tropicalis* isolates are similar to those observed in the hybrid species *C. metapsilosis* and in hybrid isolates of *C. orthopsilosis* [26–28]. This suggests that these *C. tropicalis* isolates may also be hybrids, that is, they may have at least one different parent to most *C. tropicalis* isolates. Hybrid genomes are characterized by regions of heterozygosity due to differences between the homeologous chromosomes, alternating with regions of homozygosity. This results in distinct bimodal patterns of subsequences (*k*-mers) in sequencing reads, which represent the heterozygous and homozygous regions of the genome. Such bimodal *k*-mer patterns are observed in hybrid isolates of *C. orthopsilosis, C. metapsilosis, C. inconspicua* and *C. albicans* [25,39]. We find that the *k*-mer frequency distribution of four of the six divergent *C. tropicalis* is also bimodal, with one peak at approximately 100X (the average genome-wide coverage) and one at approximately 50X (half the average genome-wide coverage) (Fig. 2A). The full and half coverage peaks represent homozygous regions and heterozygous regions respectively. Approximately half of the heterozygous *k*-mers (i.e. *k*-mers that map to heterozygous regions of the genome) are not represented in the reference genome sequence, which is a collapsed haploid reference sequence from a non-hybrid isolate (*C. tropicalis* MYA-3404). For the remaining two divergent isolates (*C. tropicalis* ct25 and ct42), the sequence coverage was too low to measure *k*-mer distribution. This analysis suggests that at least four of the divergent isolates are hybrids, resulting from mating between two related, but distinct, parents. For all four isolates, the heterozygous peak is considerably higher than the homozygous peak, indicating that the hybridization event(s) are recent, and very little loss of heterozygosity (LOH) has occurred.

**Figure 2.**
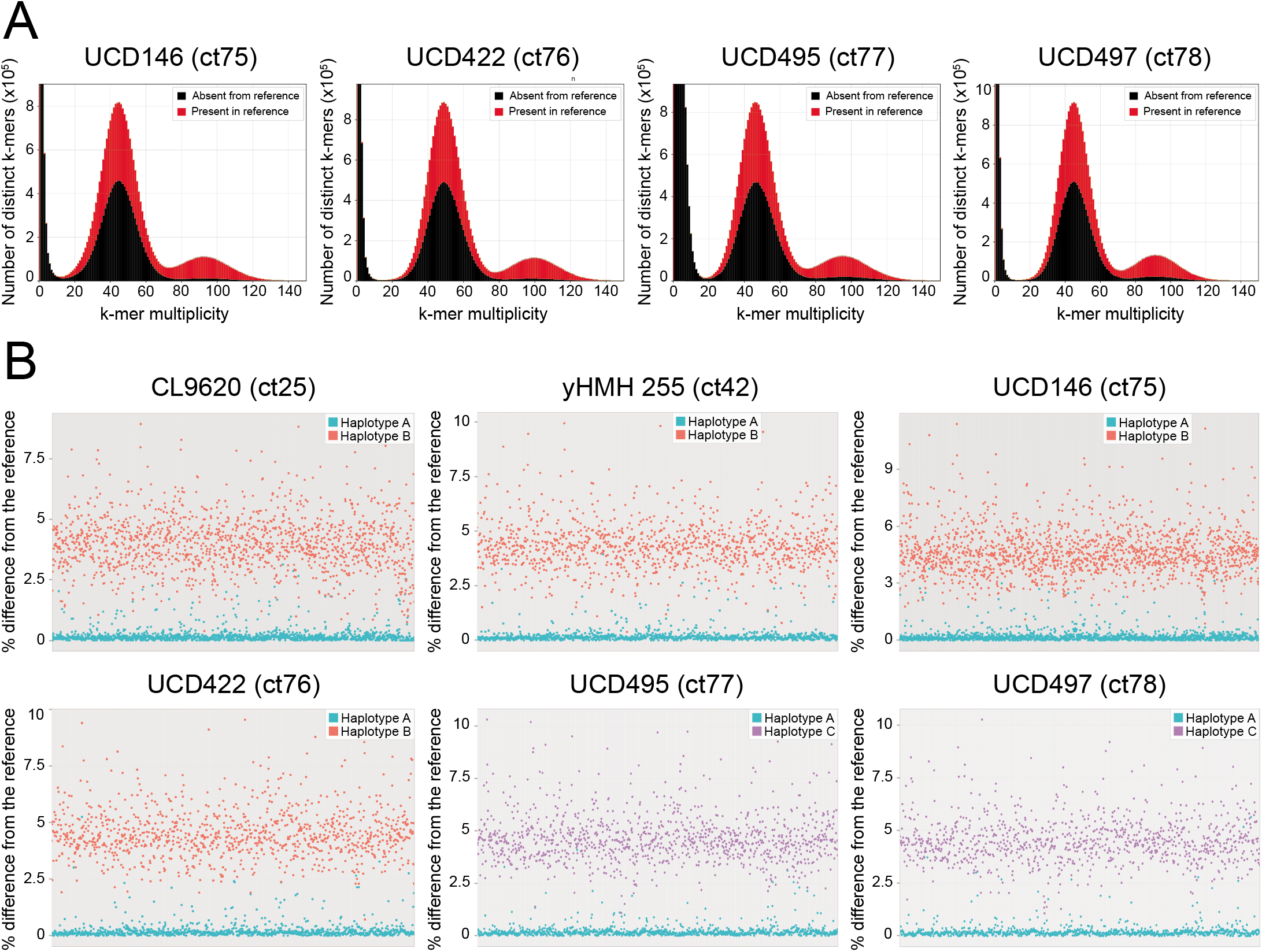
Novel *C. tropicalis* isolates result from hybridization. **(A) Analysis of *k*-mer distribution profiles reveals hybrid genomes.** *K*-mer analysis of sequencing readsets was performed with the *k*-mer Analysis Toolkit (KAT [53]). For each of four divergent isolates, the number of distinct *k*-mers of length 27 bases (27-mers) is displayed on the Y-axis and *k*-mer multiplicity (depth of coverage) is displayed on the X-axis. *K*-mers that are present in the reference genome are shown in red, and *k*-mer that are absent from the reference genome are shown in black. There are two distinct peaks of *k*-mer coverage at approximately 50X and 100X. This pattern implies that most of the genomes are heterozygous (*k*-mers at 50X coverage) with few homozygous regions (*k*-mers at 100X coverage). Approximately half of the heterozygous *k*-mers in the readsets are not represented in the reference sequence. This pattern has been observed in hybrid isolates from other yeast species [25]. **(B) Analysis of phased variants identifies two distinct haplotypes in divergent isolates of *C. tropicalis*.** Variants were phased using HapCUT2 [45] into blocks covering 10 - 12 Mb of the genome. For each phased block, percentage difference from the reference strain was calculated as the number of variants divided by the length of the block. For 84 - 87% of the blocks, one haplotype is <0.3% different to the reference sequence and one haplotype is >4% different to the reference sequence. All phased blocks for each of the six hybrid isolates are shown as pairs, with the member of the pair more similar to the reference (haplotype A) shown in blue and the member of the pair less similar to the reference shown in orange (haplotype B) or purple (haplotype C).

To further investigate the origins of the six divergent isolates, we attempted to separate the haplotypes of the two parental chromosomes. Approximately 500,000 - 700,000 heterozygous sites were identified per isolate. The heterozygous sites were placed in phased blocks, using HapCUT2 [45]. On average, 86% of the variants in each isolate were successfully phased, with a total phased span in base pairs of approximately 10 - 13 Mb (Table 1).

**Table 1.** Results of haplotype phasing.

|  |  |  |  |  |  |  |
| --- | --- | --- | --- | --- | --- | --- |
|  | CL9620 | yHMH25<br>5 | UCD146 | UCD422 | UCD495 | UCD497 |
| **Total number of heterozygous variants** | 526,189 | 638,854 | 691,443 | 707,685 | 697,033 | 685,835 |
| **Variants successfully phased** | 462,386<br>(88%) | 551,867<br>(86%) | 589,165<br>(85%) | 602,663<br>(85%) | 592,497<br>(85%) | 583,248<br>(85%) |
| **Total phased span (bp)** | 10,850,562 | 12,412,152 | 12,431,473 | 13,046,231 | 12,672,096 | 12,629,063 |

For each phased block of the genome greater than 1 kb, the percentage difference of each haplotype to the reference sequence was calculated. For the majority of blocks (84 - 87%), one haplotype (which we refer to as haplotype A) has >99.7% identity to the reference and the second haplotype is more than 4% different to the reference (Fig. 2B). The alternative haplotypes were constructed by substituting all variant sites in the reference sequence with alleles that had been assigned to the alternative haplotype. The alternative haplotypes of all six isolates are 4.0 - 4.6% different from the reference strain. The alternative haplotypes of four of these isolates, *C. tropicalis* ct25, ct42, ct75 and ct76), which we refer to as haplotype B, are approximately 1% different from each other. The alternative haplotypes of the other two, *C. tropicalis* ct77 and ct78, called haplotype C, are approximately 3% different in sequence to the B haplotypes in the other four isolates (and less than 1% different in sequence from each other).

These analyses strongly suggest that the six novel isolates originated from mating or hybridization between related parents, one of which is very similar to the *C. tropicalis* reference, and others that are > 4% different. The second parent is not the same for the six divergent isolates. We therefore refer to most *C. tropicalis* isolates as AA diploids, to four isolates as AB diploids, and to two isolates as AC diploids. All AB and AC isolates contain only one rDNA locus (D1/D2 region), which is 99% identical to the reference haplotype A. The rDNA sequences were confirmed by PCR amplification and Sanger sequencing (Supplementary file S1).

### Loss of heterozygosity (LOH) in *C. tropicalis* isolates

Loss of heterozygosity (LOH) describes tracts of the genome that are essentially homozygous, most likely due to gene conversion or mitotic recombination. We observe a pattern of heterozygous regions alternating with homozygous (LOH) regions in all *C. tropicalis* isolates (Fig. 3A). We defined heterozygous regions of the genome as regions of at least 100 bp in length containing at least two heterozygous variants; all remaining regions of the genome were classified as homozygous, or LOH, regions, as long as they were at least 100 bp in length.

**Figure 3.**
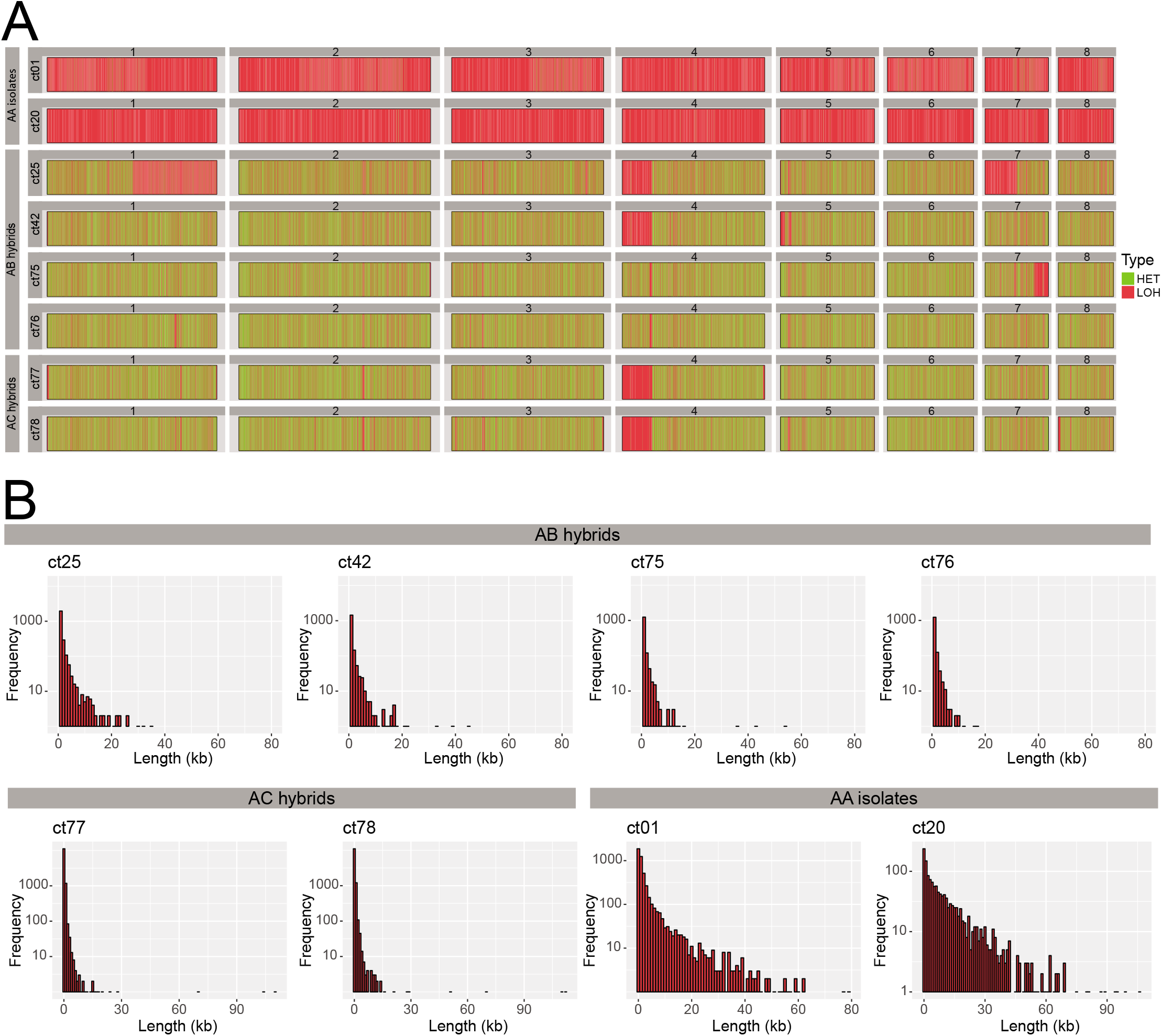
Loss of heterozygosity in *C. tropicalis* isolates. **(A) Hybrid and non-hybrid isolates differ in the extent of LOH across the genome.** The eight largest scaffolds in the reference genome are displayed horizontally from left to right and labelled from 1 to 8. LOH blocks are shown in pink and heterozygous (“HET”) blocks are shown in green. Isolates are labelled on the left-hand side. *C. tropicalis* ct01 is shown as a representative of the non-hybrid (AA) isolates. The genomes of the AA isolates consist mostly of LOH blocks. The AA isolate *C. tropicalis* ct20 has undergone extensive LOH, covering >99% of the genome. In contrast, in the AB/AC isolates, the majority of the genome consists of heterozygous blocks. **(B) LOH is limited to short tracts of the genome in hybrid isolates.** The histograms show the frequency of LOH blocks of different lengths in the six hybrid isolates and two AA (non-hybrid) isolates *C. tropicalis* ct01 and *C. tropicalis* ct20. Frequency is shown on a log scale on the Y-axis while length in base pairs (bp) is shown on the X-axis, with a bin width of 1000 bp. The average length of LOH blocks in the hybrid isolates ranges from 286 - 416 bp. A similar pattern is observed in all six hybrid isolates, i.e. a predominance of short LOH blocks, with very few long tracts of LOH. In the non-hybrid isolates (e.g. *C. tropicalis* ct01), LOH blocks are generally longer. *C. tropicalis* ct20 has the longest average LOH block length (~10 kb).

Only 4% on average of the non-hybrid (AA) genomes are heterozygous, with heterozygous blocks with a mean length of 208 bp and a maximum length of approximately 7.6 kb (Table S5A). In *C. tropicalis* ct20 only 0.37% of the genome is heterozygous, with a mean block length of 213 bp. In contrast, on average, 69% of the six hybrid genomes consists of heterozygous regions, with a mean length of approximately 900 bp, and a maximum length of approximately 13.8 kb.

Analysis of heterozygous regions in the six hybrid isolates reveals further support for the hypothesis that they originated from different hybridization events involving different parent strains (B and C). If we assume that the hybrid isolates were derived from a mating event between two parental isolates, we can expect that the heterozygous regions of the genome in the hybrid isolates should be derived equally from the two parent strains. Therefore, if two hybrids originated from hybridization between the same parental strains, the heterozygous regions of their genomes should carry the same variants. However, if two hybrids originated from hybridization between different parental strains, the variants in common heterozygous regions will be different. Shared heterozygous regions were defined as regions of heterozygosity in the hybrid isolates that share exact boundaries. Shared heterozygous regions in all six hybrid isolates cover 5.8 Mb, with only 217,997 variants (~ 45% of all heterozygous positions in these regions) present in all six. This indicates that the six hybrid isolates did not all originate from the same parental strains. However, there is a much higher degree of conservation of variants in shared heterozygous regions among the four AB isolates; 94% of 419,440 heterozygous variants in 6.7 Mb are present in all four. Similarly, the two AC hybrids share 98% of 620,569 variants across 9.6 Mb. This further indicates (in line with our previous analyses) that the four AB isolates share a common origin, and that the two AC isolates share a common origin that is separate from the origin of the AB isolates.

There is extensive LOH in the non-hybrid isolates, covering on average 95% of the genome (Table S5B). In *C. tropicalis* ct20, >99% of the genome is in LOH blocks. The average length of all LOH blocks across all non-hybrid isolates (excluding *C. tropicalis* ct20) is approximately 1.7 kb with a maximum length of 238 kb. In contrast, limited LOH is observed in the six hybrid (AB/AC) isolates, with an average of 13,139 LOH blocks of at least 100 bp, covering between 25 and 42% of the genome. The average length of LOH blocks in the AB/AC isolates is 330 bp, but can be as long as 112 kb (Fig. 3B). Only 1.6% of LOH blocks (equating to 731 LOH blocks) is conserved among all six isolates. There are more shared LOH regions in the four AB isolates; 17% of LOH blocks (equating to 5,131 LOH blocks) in these isolates are identical. In the AC isolates, 55% of LOH blocks are identical (equating to 8,807 LOH blocks). There is a large LOH block at the start of scaffold 4 (equivalent to Chromosome R [40]) covering approximately 400 kb, that is shared between four of the hybrid isolates (*C. tropicalis* ct25, ct42, ct77 and ct78). The LOH block extends from the telomere to the rDNA locus, although the exact end point differs, and it is interrupted by some small heterozygous regions. A larger LOH block, encompassing this region and extending to the centromere, was identified in a complete, chromosome-scale assembly of *C. tropicalis* and in the related species *Candida sojae* [40]. Two of the AB hybrids (*C. tropicalis* ct75 and ct76) are unique, in that only the rDNA locus itself has undergone LOH.

We considered the possibility that the homozygous isolate *C. tropicalis* ct20 might represent one parent of the hybrid isolates. We therefore compared it with both haplotype A and haplotypes B and C of the six hybrid isolates by computationally reconstructing both subgenomes of each hybrid strain. We constructed a putative A haplotype from *C. tropicalis* ct20 by substituting bases in the reference with homozygous variants identified in this isolate. For the hybrid isolates, the A haplotype was constructed by substituting variants that were originally assigned to haplotype A during haplotype phasing (see Materials & Methods, subsection Haplotype splitting). Similarly, B and C haplotypes were constructed by substituting variants that were assigned to either B or C. The A haplotypes from the hybrids share, on average, approximately 8% of variants with *C. tropicalis* ct20 (i.e. approximately 8% of variants identified in *C. tropicalis* ct20 and a given hybrid isolate are identical). There is even less similarity between the B and C haplotypes and *C. tropicalis* ct20; only 1% of variant sites in *C. tropicalis* ct20 and the hybrid haplotypes B or C are identical. *C. tropicalis* ct20 therefore has a A haplotype, but it is unlikely that it is a parent, or closely related to a parent, of the hybrid isolates.

### Mating type-like loci (MTL) in *C. tropicalis* isolates

Most AA isolates (46) are heterozygous at the *MTL*, similar to previous reports [20,21] (Table S1). In addition, two heterozygous isolates have three copies of the MTL. The triploid isolate *C. tropicalis* ct66 is *MTL* **a**/**a**/*α* (Figure S3). *C. tropicalis* ct18 is trisomic for scaffold 8, which carries the *MTL*, and is *MTL***a**/*α/α*. Fourteen are homozygous for *MTL***a**/**a** and seven are homozygous for *MTLα/α*. In addition, *C. tropicalis* ct06 is trisomic for scaffold 8, which carries the *MTL*, and has three copies of *MTLα*. The *MTL* idiomorphs of the octaploid isolate, *C. tropicalis* ct26, could not be definitively determined by assembling the Illumina data or by PCR, but it appears to have 7 copies of *MTLα* and one copy of *MTL***a** (Fig. S3).

All six AB and AC isolates contain both *MTL***a** and *MTLα* idiomorphs. In the AB isolates, the *MTL***a** idiomorphs are >99% identical to that of the reference strain (A haplotype) with only three nucleotide changes across the entire locus (8,180 bp). These include synonymous and nonsynonymous substitutions in *PAP***a** and *PIK***a**. In addition, one isolate (*C. tropicalis* ct42) has a nonsynonymous substitution in *MTL***a***1*. Apart from this, the *MTL***a** idiomorphs in the AB isolates are identical. The *MTL***a** idiomorph therefore likely originated from the A parent. The *MTLα* loci are >99% identical in all four AB isolates, and ~7% different to the reference strain, indicating that it was donated by the B parent. All AB isolates therefore most likely resulted from mating between the same parents, an *MTL***a** parent similar to the reference strain (parent A), and an *MTLα* parent which is approximately 4% different (parent B).

In the two AC isolates, the *MTLα* idiomorphs are also identical to each other, and they are >99% identical to the reference strain. *MTL***a** idiomorphs are identical to each other, and approximately 96% identical to the reference strain. The *MTL***a** idiomorph in the AC isolates therefore originated from the C parent, and the *MTLα* idiomorph originated from the A parent.

### Analysis of phenotypic variation in *C. tropicalis*

To measure the phenotypic diversity within *C. tropicalis*, the growth of 68 AA isolates was tested in 61 different conditions, including alternative carbon sources, stressors (e.g. calcofluor white, congo red), heavy metals (e.g. zinc, cobalt, cadmium) and antifungal drugs (e.g. fluconazole, ketoconazole, caspofungin) (Fig. S5A). Because nitrogen and carbon metabolism are important virulence attributes in fungi [46], the ability of *C. tropicalis* isolates to use different sole nitrogen sources (e.g. amino acids, gamma-aminobutyric acid (GABA)) was also tested (Fig. S5B). The AB and AC isolates and the engineered lab isolates *C. tropicalis* ct13, ct14 and ct15 were excluded from the analysis.

The *C. tropicalis* isolates show wide variation in their growth characteristics (Fig. S5). We attempted to identify genome variants that are associated with specific growth defects. For this analysis, only conditions that resulted in a growth defect of at least 70% compared to the control condition in at least one strain were included (i.e. 25 conditions using YPD as a base media, and 10 conditions using different nitrogen sources). Reduced growth was scored as 1, and growth similar to the control was scored as 0. Predicted genomic variants were annotated with SnpEff [47] to identify those that were likely to have a major impact on protein function. 390,321 variant sites were identified in total across 68 isolates. The majority of variants (~75%) were SNPs, with the remainder consisting of small insertions and deletions (indels) (Fig. S4A). Most variants are found in intergenic regions, or are silent or missense mutations. Only variants that were predicted to have a high impact, including frameshifts, gene fusion events, loss or gain of a stop codon, or variation at splice donor or acceptor sites (9,261 variants, Fig. S4B), were included in the genotype-phenotype correlation analysis.

One clinical isolate, *C. tropicalis* ct04, identified by cosine similarity analysis [48], has impaired growth when valine or isoleucine (branched chain amino acids) are provided as the sole nitrogen source (Fig. 4A). Compared to other isolates, *C. tropicalis* ct04 also grows poorly on 2% sodium acetate, 2% starch and in the absence of a carbon source. There are 40 variants unique to this isolate that are predicted to have a high impact on protein function (Table S6). One of these is a heterozygous deletion of two bases in CTRG_06204 (*BAT22*), an orthologue of the *S. cerevisiae BAT1/2* genes that encode a branched-chain amino acid aminotransferase (BCAT). BCATs catalyze the final step of biosynthesis and the first step in the degradation of the branched chain amino acids valine, isoleucine and leucine [49]. The deletion results in a frameshift which introduces a premature stop codon at amino acid Gly30 of the Bat22 protein (Fig. 4B). We determined if introducing an equivalent change into other genetic backgrounds using CRISPR/Cas9 [50] would result in the same phenotype. A repair template was designed to delete two bases and also to destroy the target of the guide RNA to prevent recutting. The gene was edited in three different *C. tropicalis* isolates ct09, ct44 and ct53. All edited strains can no longer use valine or isoleucine as sole nitrogen sources (Fig. 4C). However, unlike *C. tropicalis* ct04 they have no growth defect on sodium acetate, starch or in the absence of carbon sources, indicating that another variant, or combination of variants, is responsible for these phenotypes.

**Figure 4.**
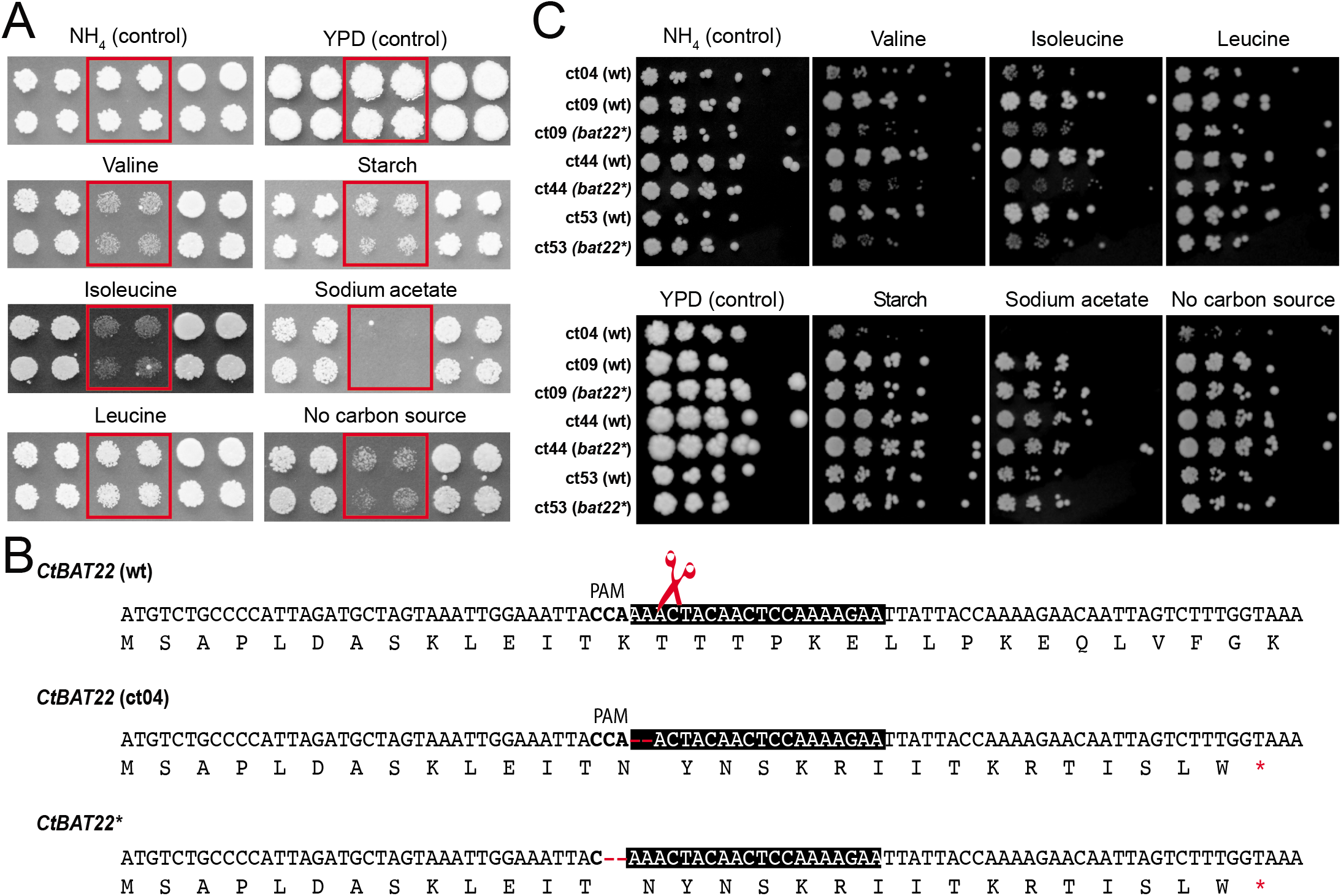
Disrupting *BAT22* prevents growth of *C. tropicalis* on branched chain amino acids as a sole nitrogen source. **(A)** Growth of *C. tropicalis* ct04 is shown on solid media. Strains were grown in 2×2 arrays; two biological replicates (top and bottom rows), with two technical replicates each (left and right columns), of each strain were tested. *C. tropicalis* ct04 replicates are outlined with red boxes. *C. tropicalis* ct04 cannot utilize valine or isoleucine as a sole nitrogen source and also exhibits a growth defect on solid media with 2% starch or 2% sodium acetate as the sole carbon source, or on solid media without a carbon source provided. **(B)** Plasmid pCT-tRNA-BAT22 was generated to edit the wild type sequence of *BAT22* (*CTRG_06204*) using CRISPR-Cas9. The sequences of the reference *C. tropicalis BAT22 (CtBAT22* (wt)), *BAT22* from *C. tropicalis* ct04 (*CtBAT22* (ct04)) and edited *BAT22* (*CtBAT22\**) are shown. The guide sequence is highlighted with a black box, the PAM sequence is shown in bold, and the Cas9 cut site is indicated with a red scissors. *C. tropicalis* isolates ct44, ct09 and ct53 were transformed with pCT-tRNA-BAT22 and a repair template (RT_BAT22_2bpDel_SNP) generated by overlapping PCR using RT_BAT22_2bpDel_SNP-TOP/BOT oligonucleotides. The repair template contains two 60 bp homology arms and deletes two bases in *BAT22* resulting in the same frameshift observed in *C. tropicalis* ct04. **(C)** 5-fold serial dilutions of *C. tropicalis* ct04, ct09(wt; bat22**), ct44 (wt; bat22**) and ct53 (wt; bat22**) in the same conditions tested in (A). The edited strains cannot use valine or isoleucine as sole nitrogen sources.

## Materials & Methods

### Strain collection and growth

*C. tropicalis* isolates were collected from a variety of clinical and environmental sources (Table S1). For phenotype analysis, isolates were inoculated as 2×2 arrays (two independent cultures with one technical replicate of each) into 200 μl of YPD broth (1% yeast extract, 2% peptone, 2% glucose) in 96-well plates and incubated at 30°C for 24 h. Stocks were diluted in 96-well plates containing 200 μl of water by dipping a 12×8 pin bolt replicator (V&P Scientific) three times in the culture and then transferring it to the water. Once diluted, the cultures were pinned onto 85 unique media on solid agar plates and incubated at 30°C for 48 h (Table S2). For 60 conditions, the base media was YPD, with 2% agar including 2% glucose as a carbon source. Glucose was substituted with different carbon sources where indicated, or compounds were added at the indicated concentrations (Table S2). To test the ability to use specific nitrogen sources (24 conditions), the base media was 0.19% of YNB (Yeast Nitrogen Base) without ammonium sulfate or amino acids, 2% glucose and 2% agar. Nitrogen sources were added as indicated (Table S2). Spider media was tested as the 85th condition (Table S2). Plates were photographed and growth was measured using SGAtools [51]. SGAtools was designed to analyze synthetic genetic interactions and assumes that average growth on a plate does not vary. This was not true for several media, where many strains grew poorly. We therefore compared the growth of each strain on the test media to the growth of the same strain on YPD, or on YNB with ammonium sulfate, as a control, using the raw data extracted from SGAtools. For each strain in each analyzed growth condition, the SGAtools scores (ranging from 0 to 1.8) were converted to a binary score where a growth ratio above 0.3 (no growth defect) was assigned 0, and a ratio below or equal to 0.3 (major growth defect) was assigned 1. These scores were chosen to be very stringent - only conditions which resulted in reducing growth to approximately 30% of that under the control conditions were judged as a defect. We found that SGAtools could not reproducibly identify enhanced growth in these conditions. The raw data for the image analysis is available at https://figshare.com/s/e0bbb5fc9e92bfd878f2.

### Genome sequencing

For most *C. tropicalis* isolates, genomic DNA was isolated by phenol-chloroform extraction followed by purification using the Genomic DNA Cleanup and Concentration kit from Zymo Research (catalogue number D4065). For three isolates (*C. tropicalis* ct76, ct77 and ct78), genomic DNA was extracted and purified using the QIAamp DNA Mini Kit from QIAGEN (catalogue number 51304). For most isolates, library preparation and sequencing was performed at the Earlham Institute, Norwich, UK using the LITE method (Low Input Transposase-Enabled), a custom Nextera-based system. These isolates were sequenced on two lanes of an Illumina HiSeq 2500 generating 2×250 bp paired-end reads. For five isolates (*C. tropicalis* ct51, ct75, ct76, ct77 and ct78), library preparation and sequencing was performed by BGI, Hong Kong, generating 2×150 bp paired-end reads, on an Illumina HiSeq 4000. Our genome sequences of two isolates (*C. tropicalis* ct20 and ct21) were almost identical. These may represent independent isolates of the same strain, or one isolate may have been accidentally sequenced twice. We therefore included only one of these (*C. tropicalis* ct20) in subsequent analysis.

For the 72 unique isolates sequenced using the LITE method, Nextera adapters were removed using TrimGalore v0.4.3 with the parameters “--paired” “--length 35” “--nextera” and “--stringency 3”. Custom adapters and low-quality bases were trimmed using Skewer v0.2.2 with the parameters “-m pe” “-l 35” “-q 30” “-Q 30” [52]. For 5 isolates sequenced by BGI, adapters were removed by the sequencing provider and reads were quality trimmed using Skewer. *K*-mer distribution profiles were analysed using the *k*-mer Analysis Toolkit v2.4.2 using the default *k*-mer length of 27 bases [53]. All genomes were assembled using SPAdes v3.9.1 with parameters “--careful” “-t 12” “-m 60” [54]. Assembly statistics were assessed using QUAST v4.4 [55]. To confirm the species identity of hybrid isolates, the D1/D2 domain of the large subunit of the ribosomal DNA was amplified using standard universal primers NL-1 and NL-4 (Table S3).

### Mating type-like locus analysis

The *MTL* idiomorph of a subset of isolates was confirmed by PCR using primer pairs *MTL***a***1*F and *MTL***a***1*R to amplify the *MTL***a***1* gene and *MTLα2F* and *MTLα2R* to amplify the *MTLα2* gene, as described in Xie et al. [21]. Colony PCR was performed by boiling single colonies in 5 μl sterile deionized water, then adding 12.5 μl MyTaq Red Mix (2X), 1 μl forward primer (100 μM), 1 μl reverse primer (100 μM) and 5.5 μl deionized water. PCR was run for 1 min at 95°C; then for 30 cycles of 30 sec at 95°C, 30 sec at 57°C, 60 sec at 72°C; and then a final 2 min at 72°C.

### *C. tropicalis* reference genome

The *C. tropicalis* reference genome annotation was updated using RNAseq data for three *C. tropicalis* strains downloaded from NCBI under BioProject ID PRJNA290183 [56]. RNAseq data were aligned against the original *C. tropicalis* reference [16] with HISAT2 v2.0.5 with the parameter “-- novel-splicesite-outfile” to predict splice sites in the genome [57]. Predicted splice sites were manually validated by examination of transcripts mapping to predicted splice sites. The reference genome sequence was subsequently scaffolded from 23 supercontigs to 16 supercontigs. Areas of overlap between supercontigs in the original reference assembly were identified using Gepard to generate dot matrix plots [58]. Overlapping supercontigs were merged if this arrangement was supported by synteny with other *Candida* species, using the *Candida* Gene Order Browser (CGOB) [59], and by data from Illumina resequencing of the reference strain. The final assembly (also known as Assembly B [40]) contained 16 supercontigs and is available at https://figshare.com/s/e0bbb5fc9e92bfd878f2. The *C. tropicalis* reference was subsequently further improved as described by Guin et al [40].

### Variant calling

For isolates sequenced using the LITE method, trimmed reads were aligned to *C. tropicalis* MYA-3404 Assembly B with bwa mem v0.7.11 to generate two BAM files per sample (one for each lane used for sequencing) [60]. BAM files were sorted with SAMtools v1.7 [61], and duplicate reads were marked using GenomeAnalysisToolkit (GATK) v3.7 Mark Duplicates [43]. BAM files from separate lanes were combined for each sample and marked for duplicates again using GATK MarkDuplicates. For isolates sequenced at BGI, Hong Kong, trimmed reads were aligned to the updated *C. tropicalis* MYA-3404 Assembly B with bwa mem v0.7.11 as before, generating only one BAM file per sample (each of these samples was sequenced on only one lane of the sequencer). BAM files were sorted with SAMtools v1.7 [61] and duplicate reads were marked using GenomeAnalysisToolkit (GATK) version 3.7 Mark Duplicates [43].

The subsequent steps were applied to all samples. Realignment around indel sites was performed using GATK IndelRealigner and variants were called using GATK HaplotypeCaller in “--genotyping_mode DISCOVERY”. Variants were filtered for quality based on genotype quality (GQ) < 20 and read depth (DP) < 10. For SNP trees, gVCFs were generated using GATK HaplotypeCaller with the parameters “-- genotyping_mode DISCOVERY” and “--emitRefConfidence GVCF”. Joint genotyping was performed using GATK GenotypeGVCFs to produce a single multi-sample gVCF. SNPs were extracted from the multi-sample gVCF using GATK SelectVariants with parameter “-selectType SNP”. Variants were filtered based on genotype quality (GQ) < 20 and read depth (DP) < 10. For genotype-phenotype analysis, the presence of a variant at a particular site in each isolate was scored as 1, and absence was scored as 0.

### Aneuploidy analysis

To calculate copy number variants based on coverage discrepancies, the *C. tropicalis* MYA-3404 Assembly B genome was split into 1 kilobase (kb) windows using the “makewindows” command from bedtools v2.26.0, with parameters “-i winnum” (label windows sequentially) “-w 1000” (window size 1 kb) [62]. Mean coverage in each 1 kb window was calculated for each sample using the “coverage” command from bedtools [62]. Average whole genome coverage for each strain was calculated using GATK DepthOfCoverage [43]. Coverage ratios for each 1 kb window were calculated as log2(window coverage / average whole genome coverage). A value of zero was assigned to windows that had zero coverage. The resultant ratios were visualized using the DNACopy package from Bioconductor in R [63]. Ploidy was also visualized using allele frequencies from heterozygous biallelic SNPs extracted from the VCF files using GATK SelectVariants with parameters “-selectType SNP” and “-restrictAllelesTo BIALLELIC”. Allele frequency was calculated as allele depth (AD) / read depth (DP). Histograms of allele frequency for each scaffold in each sample were visualized in R using ggplot2 [64].

### Phylogeny

SNP trees were drawn from filtered variants, using only those SNPs that passed the filters described in “Variant Calling”. To account for heterozygous SNPs, the Repeated Random Haplotype Sampling tool (RRHS) v1.0.0.2 was used to select a random allele at heterozygous SNP sites [65]. This process was performed 100 times to generate 100 SNP profiles for each isolate, thereby encapsulating the full heterozygosity of each isolate. For homozygous variant sites, the alternate allele was chosen by default. 100 maximum likelihood (ML) trees were drawn (one for each SNP profile) using RAxML v8.2.12 [66] with the “GTRGAMMA” model. The best-scoring ML tree was chosen as a reference tree and the remaining 99 ML trees were used as pseudo-bootstrap trees to generate a supertree using RAxML v8.2.12 with options “-f b” (draw bipartition information on a reference tree based on multiple trees (e.g. from a bootstrap)) and the “GTRGAMMA” model. Phylogeny was also examined using principal component analysis (PCA) with the ade4 package in R [67].

### Loss of heterozygosity

Loss of heterozygosity (LOH) was calculated in blocks of at least 100 base pairs (bp) across the genome. Heterozygous regions were defined as any region containing at least two heterozygous variants within 100 bp of each other, with a minimum total length of 100 bp. Remaining regions were defined as homozygous, or LOH, regions as long as they were at least 100 bp in length. Heterozygous regions shared by all isolates were identified using bedops intersect [68]. In the case of heterozygous regions that were partially shared, the portion that was common to all isolates was extracted and analysed as a shared heterozygous region. The number of common variants in the shared heterozygous regions was counted as the number of variant sites in these regions with the same genotype in all isolates. Shared LOH regions were defined as LOH blocks with identical start and stop coordinates in the relevant isolates.

### Haplotype splitting

Hybrid haplotypes were phased using HapCUT2 v0.7 [45]. The filtered variants were used as input for the subcommand “extractHAIRS” (extract haplotype-informative reads) to identify “haplotype-informative reads”, i.e. sets of reads that align to the same location in the reference genome but that contain one or more variant alleles. HapCUT2 was subsequently used to build haplotype blocks from the haplotype-informative reads with parameter threshold 30” (Phred-scaled threshold for pruning low-confidence SNPs). The difference of each phased block to the reference genome was calculated as the number of SNPs in block / length of block. Blocks were assigned to either the reference haplotype or the alternate haplotype according to their percentage difference; < 0.3% difference was assigned to reference haplotype (haplotype A) and > 1% difference was assigned to alternate haplotype (haplotype B).

### Analysis of genotype-phenotype correlation

Variants from non-hybrid isolates were further annotated with SnpEff v4.3t to predict the functional effect of variants [47]. High-impact variants (e.g. variation at splice donor or acceptor sites, variants resulting in a gain or loss of stop or start codon, or frameshifts in genes) were extracted and correlated with phenotypes. Variants were converted to binary scores; 1 for the presence of a variant in a given strain, 0 for the absence. Phenotype scores were coded as 1 for a growth defect (score of 0.3 or less), and as 0 for no growth defect (score above 0.3). For each variant-condition pair, two vectors were generated using the binary scores; the first consists of the scores for every strain with respect to the variant, the second consists of the scores for every strain with respect to the condition. For every variant-condition vector pair, the cosine similarity between the two vectors was calculated as *cos θ* = *a*⍰. *b*⍰⍰∥ *a*⍰∥∥ *b*⍰∥⍰. Any variant-condition pair with a cosine similarity of > 0.85 was selected for further analysis.

### Editing *BAT22* with CRISPR-Cas9

A 20 bp sequence (guide RNA) targeting *C. tropicalis BAT22* (*CTRG_06204*) was designed using the web tool ChopChop [69]. The guide RNA was generated by annealing of two short oligos (g60BAT22_TOP/BOT, Table S3), and then cloned into the SapI-digested pCT-tRNA plasmid to generate plasmid pCT-tRNA-BAT22, as previously described in [50]. The repair template carrying the desired modification, including the disruption of the PAM sequence, was generated by primer extension (RT_BAT22_2bpDel_SNP-TOP/BOT) using ExTaq DNA polymerase (Takara Bio, USA). *C. tropicalis* isolates ct09, ct44 and ct53 were transformed with 5 μg pCT-tRNA-BAT22 and 25 μl of unpurified RT-BAT22_2bpDel_SNP using a previously described method [50]. Transformants were selected on YPD agar plates containing 200 μg/ml nourseothricin (NTC), incubated at 30°C for 48 h. The relevant region was amplified by PCR from two NTC-resistant transformants for each strain using primers bat22_fwd_01/bat22_rev_01 and sequenced using Sanger sequencing. The pCP-tRNA-BAT22 plasmid was cured by growing the cells in the absence of selection on YPD until they failed to grow in the presence of NTC.

### Data availability

All sequencing data is available at NCBI under BioProject accession PRJNA604451. Other data sets (i.e. *C. tropicalis* genome assembly, variant calls and images for phenotype analysis) is available at https://figshare.com/s/e0bbb5fc9e92bfd878f2.

## Discussion

Like many opportunistic pathogens of humans, the natural habitat of *C. tropicalis* is unclear. Although *C. tropicalis* is well-adapted to humans, isolates are also commonly isolated from a variety of sources, including soil, sand, animal feces, by-products of industrial food production and the surface of fruits [70–75]. *C. tropicalis* is also a component of the human oral and gastrointestinal mycobiome [76,77] and has been isolated from human skin [78] and the gastrointestinal tracts of mice [79].

Enrichment of *C. tropicalis* in the gastrointestinal tract has been associated with Crohn’s disease, potentially due to its invasive abilities [77].

We found little evidence of clade structure associated with geographical origin, suggesting that there may be a high degree of admixture between *C. tropicalis* populations from different regions. This is similar to what has been observed in other diploid CUG-Ser1 clade species, e.g. *C. metapsilosis* [38], *C. orthopsilosis* [28] and *C. albicans*, other than the “*C. africana”* lineage [24]. Some studies have suggested that population structure in the bakers’ yeast *S. cerevisiae* is more related to ecological niche than to geography [80,81], while others found no clear separation between different ecological groups, such as pathogenic and non-pathogenic isolates [82].

Mixao et al [25] suggested that *C. tropicalis* isolates are standard diploids, i.e that the two parents were closely related. In contrast, *C. metapsilosis* and *C. albicans* isolates descended from ancient hybridizations between two related parents, and hybridization in *C. orthopsilosis* is ongoing [25,26,28,38]. We have now shown that six divergent isolates of *C. tropicalis* result from hybridization between one parent that is highly similar in its sequence to the reference genome (parental haplotype A), and other unidentified parents (parental haplotype B or C) that are approximately 4% different in sequence to the reference strain. The low level of LOH in the *C. tropicalis* AB and AC isolates suggests that hybridization has occurred relatively recently. In addition, the isolation of hybrids from different geographical locations, and the identification of multiple hybrids originating from separate hybridization events, indicates that hybridization may be ongoing in this species. This contrasts with *C. albicans* and *C. metapsilosis*, where it is proposed that all known isolates originated from a single hybridization event [25,38], and *C. orthopsilosis*, where several hybridizations have occurred but there has been substantial LOH [28]. In addition, we identified one highly homozygous AA isolate (*C. tropicalis* ct20). This may have resulted from major loss of homozygosity in a non-hybrid isolate, similar to that proposed for the *C. africana* lineage [25]. It is also possible that homozygous isolates are the parents of hybrid isolates that have not yet been identified.

Ongoing hybridization has been associated with virulence in both plant and animal fungal pathogens [83,84]. In particular, hybridization has been proposed to facilitate the emergence of virulence in species within the CUG-Ser1 clade [85], based on the observation that most isolates of *C. albicans, C. orthopsilosis* and *C. metapsilosis* are hybrids [25,26,28,38,85]. In addition, clinical isolates of *S. cerevisiae* are more heterozygous than non-clinical isolates, indicating that heterozygous isolates may have an advantage in the human host environment [82]. However, we found that *C. tropicalis* hybrids are rare (6 of 77 isolates), and only one of these was from a clinical setting. In contrast, five of twelve environmental isolates were hybrids, suggesting that hybridization may be advantageous in non-clinical settings. The hybrid isolates we identified are heterozygous at the mating-type like locus, suggesting that they originated by mating [17].

The definition of species is a challenging and controversial topic in biology, particularly so in the case of microorganisms [86]. The level of divergence that we observe between the A and B/C haplotypes in the *C. tropicalis* hybrids is greater than the level of divergence generally observed between strains of the same yeast species. For example, the maximum divergence between strains of *S. cerevisiae* is 1.1% [87], although the divergence between distant isolates of *Saccharomyces paradoxus* or *S. kudriavzevii* can be as high as 4.6% [86]. However, high levels of divergence between parents can be tolerated during hybridization. For example, the parents of the hybrid *M. sorbitophila* are estimated to diverge by approximately 11% [29]. It is clear that species definition in fungi, and in particular in CUG-Ser1 clade yeasts, needs to include hybridization [85]. It has been suggested that the *C. parapsilosis* clade (which currently consists of three species; *C. parapsilosis* sensu stricto, *C. orthopsilosis* and *C. metapsilosis*) should be reorganized to include homozygous lineages (of which there are at least five) and heterozygous lineages (of which there are at least two) [38]. Several of the proposed homozygous lineages are uncharacterized, or only partially characterized. We have shown that *C. tropicalis* isolates can be subdivided into at least three groups; the AA lineage (where either A haplotype may carry the *MTL***a** or *MTLα* idiomorph), the AB lineage (with *MTL***a** from the A haplotype) and the AC lineage (with *MTLα* from the A haplotype). The majority of AA isolates retain some heterozygosity, including at *MTL*. However, one AA isolate (*C. tropicalis* ct20, *MTL***a**/**a**), which may have undergone extensive LOH, has approximately one heterozygous variant every 1,190 bases. This is similar to *C. dubliniensis* (approximately one SNP every 1,511 bases [88]), but not quite as homozygous as *C. parapsilosis* (on average, one SNP per 15,553 bases [16]) or homozygous isolates of *C. orthopsilosis* (approximately one heterozygous SNP per 10,692 bases [26]. Further work is required to fully characterize the individual haplotypes of each lineage. For example, long-read sequencing may be useful to produce complete, phased diploid genome sequences of each lineage.

We attempted to correlate genetic variants with phenotypes in the *C. tropicalis* AA isolates. Previous studies using MLST suggested that certain characteristics may be clade-specific in *C. tropicalis*, e.g. increased resistance to antifungals including fluconazole and flucytosine [23,89,90]. There are several difficulties with using genome-wide association studies (GWAS) to identify causative variants in fungi, including small sample sizes (in comparison to human studies), structural variation between isolates, and the influence of population structure [91,92]. In addition, phenotypes are often caused by a complex network of genetic and environmental factors. However, we previously applied cosine similarity to identify phenotype-genotype correlations in the related species *C. orthopsilosis* [48], by converting variants and phenotypes in different growth conditions to binary scores (presence/absence). A similar analysis allowed us to identify a variant in *BAT22* in one *C. tropicalis* isolate that is associated with the inability to use valine or isoleucine as sole nitrogen sources. However, the method has its drawbacks. For example, *C. tropicalis* ct04 has defects in many growth conditions other than valine or isoleucine, and contains at least 40 variants with respect to the reference strain with predicted high impact. The *BAT22* variant was selected based on information available from orthologs in *S. cerevisiae* and *C. albicans*.

*S. cerevisiae* encodes two BCAT enzymes, Bat1p (found in the mitochondria) and Bat2p (found in the cytosol) [93,94]. *BAT2* is mainly associated with catabolism and *BAT1* with biosynthesis of the branched chain amino acids valine, isoleucine and leucine [49,95,96]. Many *Candida* species, including *C. tropicalis*, also have two BCAT isozymes, which result from a recent gene duplication event [97]. *C. tropicalis* ct04 (*bat22*) has growth defects when either valine or isoleucine are the sole nitrogen source, but not when leucine is the sole nitrogen source. Previous studies have shown that leucine metabolism can occur in *S. cerevisiae* even when BCATs are deleted [49,96]. It has therefore been suggested that there are other unknown transaminases that contribute to leucine metabolism [49,96]. It is possible that in *C. tropicalis* catabolism of leucine requires Bat21 rather than Bat22, or other unknown transaminases.

Our study greatly expands the analyses of genotype and phenotype of *C. tropicalis* isolates. We have described the existence of hybrids for the first time in this species, and we question the hypothesis that hybridization is generally associated with virulence in CUG-Ser1 species. In addition, we have shown that genotype and phenotype correlations can be used to identify causative variants in *C. tropicalis*.

## Acknowledgements

We are grateful to Dr Shawn R. Lockhart from the Mycotic Diseases Branch, Centers for Disease Control and Prevention, United States for providing isolates. Thanks to Elizabeth Boyd, Eric Butler, Jane Kennedy and Aaron McLaughlin for collecting and identifying *C. tropicalis* isolates from soil as part of an undergraduate project at University College Dublin; Quinn K. Langdon and Dana A. Opulente for helping mentor MABH; and the Zasadil Family for collecting samples in the USA. This work was supported by grants to GB from European Union’s Horizon 2020 research and innovation program under the Marie Sklodowska-Curie grant agreement no. H2020-MSCA-ITN-2014-642095, the Wellcome Trust (grant number 109167/Z/15/Z), and Science Foundation Ireland (www.sfi.ie; 19/FFP/6668). The work from the CTH lab supported by the National Science Foundation under Grant No. DEB-1442148, in part by the DOE Great Lakes Bioenergy Research Center (DOE BER Office of Science DE-SC0018409), and the USDA National Institute of Food and Agriculture (Hatch Project 1020204). CTH is a Pew Scholar in the Biomedical Sciences and a H. I. Romnes Faculty Fellow, supported by the Pew Charitable Trusts and Office of the Vice Chancellor for Research and Graduate Education with funding from the Wisconsin Alumni Research Foundation, respectively.

## Supplementary material

**File S1. rDNA sequencing results from six hybrid *C. tropicalis* isolates.**

## Supplementary Figures

**Supplementary Figure 1. Polyploidy and aneuploidy in *C. tropicalis* isolates.**

**(A) Polyploidy of *C. tropicalis* isolates.** The frequency of the non-reference allele for all heterozygous biallelic SNPs across all scaffolds is shown for each of the isolates, with frequency on the Y-axis and alternate (non-reference) allele frequency on the X-axis. For each SNP, allele frequency was calculated as the depth of the alternate allele divided by the total depth at the variant site. Triploidy of *C. tropicalis* ct66 is indicated by peaks of allele frequency at 0.33 and 0.66. Octaploidy of *C. tropicalis* ct26 is indicated by peaks of allele frequency at approximately 0.5, 0.12 and 0.87. Allele frequencies of approximately 0.125 and 0.875 imply that seven chromosomes carry one allele, and one chromosome carries a second allele. In this isolate, we also observe a peak at 0.5, implying that in some cases, four chromosomes carry one allele and four scaffolds carry a second allele. This multimodal distribution (i.e. peaks at 0.125, 0.50 and 0.875) is likely to be the result of loss of heterozygosity (LOH) affecting portions of some scaffolds, leading to a pattern wherein some variant sites have a 4:4 ratio of reference:non-reference allele frequency and some have a 7:1 ratio.

**(B) Aneuploidy of *C. tropicalis* isolates.** Single chromosome aneuploidies were identified in three isolates; *C. tropicalis* ct06, a clinical isolate from Dublin, Ireland, *C. tropicalis* ct15, an engineered strain from the USA [42], and *C. tropicalis* ct18, a clinical isolate from Madrid, Spain. Aneuploidies were identified by patterns in the distribution of allele frequency in heterozygous biallelic SNPs (shown as red histograms for the relevant scaffold, with frequency on the Y-axis and alternative allele frequency on the X-axis). Allele frequency was calculated as the depth of coverage of the alternate (non-reference) allele divided by the total depth at the variant site. Aneuploidies were confirmed by elevated coverage at the relevant locus (shown as dot plots, with green and black representing alternating scaffolds). Scaffolds are listed in decreasing order of size; the eight largest scaffolds are shown. The equivalent chromosomes in the assembly described by Guin et al. [40] are: scaffold 1 and chromosome 3; scaffold 2 and chromosome 1; scaffold 3 and chromosome 4; scaffold 4 and chromosome R; scaffolds 5 and 6 and chromosome 2, scaffold 7 and chromosome 6; and scaffold 8 and chromosome 5.

**Supplementary Figure 2. PCA analysis of *C. tropicalis* genomes.**

Principal component analysis (PCA) of Cluster A isolates (Fig. 1) was performed using the ade4 package in R [67] (Table S4). Principal components 1 and 2 are represented on the X- and Y-axes respectively. Six clusters were identified using Ward’s method. Clusters one, three, four, five and six are the same as groupings as Fig. 1C, except that *C. tropicalis* ct09 is included in Cluster 4 in the PCA analysis only, and C. tropicalis ct38 and *C. tropicalis* ct66 are included in Cluster 1 in the PCA analysis only.

**Supplementary Figure 3. Analysis of *MTL* idiomorphs.**

The gel shows the results of the colony PCR amplification of the *MTL* in eleven *C. tropicalis* isolates (labelled in grey or white boxes). Hyperladder is shown on the left- and right-most column of the gel on both rows, with the sizes of the bottom three markers (200 bp, 400 bp and 600 bp) marked. Two reactions were performed for each isolate - one using primer pairs *MTL***a***1*F and *MTL***a***1*R to amplify the *MTL***a***1* gene (lane marked “a”) and *MTLα2F* and *MTLα2R* to amplify the *MTLα2* gene (lane marked “α”), as described in Xie et al. [21]. A band of 253 bp is expected in the “a” lane for isolates with at least one copy of the *MTL***a***1* gene and a band of 525 bp is expected in the “α” lane for isolates with at least one copy of the *MTLα2* gene. Negative control (all components of PCR mix excluding input DNA) is marked as “NC” on the bottom row, with one lane for each primer set (marked “a” and “α”). Most isolates are heterozygous, but *C. tropicalis* ct14 and ct73 are homozygous for *MTL***a**. The octoploid isolate *C. tropicalis* ct26 has a strong positive signal for *MTLα* (lane marked “α”) and a weak positive signal for *MTL***a** (lane marked “a”), highlighted with a red box. The genome assembly contains one full copy of *OBP**a***, and partial copies of the remainder of the *MTL***a** genes (*PAP***a**, *PIK***a**, *MTL***a***2* and *MTL***a***1*). The five *MTL***a** genes are scattered across five low-coverage contigs (coverage 1.3X - 2X), most of which are only the length of the gene itself. One gene, *MTL***a***2*, is split across two scaffolds. It is possible that there is one copy of *MTL***a** and up to seven copies of *MTLa*, resulting in low sequencing coverage of the *MTL***a** locus.

**Supplementary Figure 4. Variants in *C. tropicalis* isolates by category.**

**(A)** The majority of variants in non-hybrid (AA) *C. tropicalis* isolates are single nucleotide polymorphisms (SNPs). Variants were called in all non-hybrid isolates using the Genome Analysis Toolkit [43] and annotated with SnpEff [47]. Variant type is shown as a barplot, with variant categories on the X-axis and variant count on the Y-axis. Approximately 75% of all annotated variants are SNPs, 12.51% are insertions and 12.57% are deletions.

**(B)** 9,261 high-impact variants were identified across 68 non-hybrid *C. tropicalis* isolates. Variant classification according to SnpEff is shown as a barplot, with estimated impact level categories on the X-axis and variant count on the Y-axis. Precise counts are shown above each bar. 9,261 variants were annotated as “high impact.” These variants are predicted to have a major impact on protein function (e.g. gain or loss of start or stop codon, frameshifts, or splice site variants). These variants were analysed for potential genotype-phenotype correlations.

**Supplementary Figure 5. Phenotypic analysis of *C. tropicalis* AA isolates.**

68 *C. tropicalis* isolates were grown on YPD (A) or YNB with ammonium (NH4) (B) solid agar media as a control, and compared to strains growing on solid agar media containing different stressors. Pictures were taken after 48 hours and colony size and growth scores were measured using SGAtools [51]. Heatmaps show the normalized raw colony size in various tested growth conditions. Isolates are represented in rows, and are ordered alphabetically by strain alias. Growth conditions are shown in columns. Increased growth relative to YPD or YNB + NH4 is shown in green (1 - 2) and decreased growth is shown in purple (0 - 1). Major differences are observed between isolates growing in the presence of cell wall stressors (calcofluor white, congo red, sodium dodecyl sulphate, caffeine), and antifungal drugs (ketoconazole, caspofungin, fluconazole). Hybrid isolates and engineered lab isolates were excluded from this analysis.

**Supplementary tables**

**Table S1. List of strains used in this study.**

**Table S2. List of media used for phenotypic testing.**

**Table S3. List of primers used in this study.**

**Table S4. Isolate clusters identified by principal component analysis.**

**Table S5. Summary of LOH and heterozygous blocks in *C. tropicalis* isolates.**

**Table S6. List of phenotype-genotype correlations**

